# β-Catenin Drives Apical–Basal Polarization to facilitate TE lineage commitment *In Vitro* and *In Vivo*

**DOI:** 10.64898/2026.06.03.729846

**Authors:** Jie Hu, Xuzhao Zhai, Yating Zhu, Boyan Huang, Shu Sun, Jiali Fu, Wenwen Shi, Ling Li, Dan Liang, Wakam Chang, Man Zhang

## Abstract

Embryo polarization is critical for the first cell fate segregation. While mechanisms underlying its initiation have been described, the intrinsic signaling pathways that regulate this process remain poorly understood. Here, we show that mouse embryonic stem cells, when aggregated under defined conditions, recapitulate the first lineage segregation to generate trophectoderm (TE)-like cell populations and undergo self-organized morphogenesis into blastocyst-like structures. In the blastoid-forming medium, we identify CHIR99021 is essential for the generation of blastoids from both ESCs and totipotent-like cells. CHIR99021 promotes cell polarization and TE differentiation by activating the WNT/β-catenin pathway and upregulating associated genes. Consistent with this, genetic ablation of β-catenin abolished the cell polarization and disrupted blastoid formation from ESCs, a defect that was restored by β-catenin overexpression. Moreover, β-catenin depletion compromised cell polarization in natural embryos. Collectively, this study establishes the Wnt/β-catenin as a critical regulator initiating polarization *in vitro* and in mouse early embryo development.

## Introduction

In mouse early embryo development, 8-cell embryo undergoes the first cell fate decision to produce two cell types: trophectoderm (TE) and inner cell mass (ICM) ^1,2^, a process orchestrated by embryonic polarization ^3,4^. Embryonic polarization closely resembles to epithelial cell polarization, with blastomeres undergoing apical-basal polarization: the apical domain faces the external environment while the basolateral domain contacts adjacent cells ^5^. During the initial stage of polarization, phospholipase C (PLC)-mediated hydrolysis of phosphatidylinositol 4,5-bisphosphate (PIP2) activates protein kinase C (PKC), and in turn RhoA, leading to cortical accumulation of actomyosin ^6,7^. This actomyosin polarization is a prerequisite for the second phase, in which other polarity proteins including the PAR complex (PAR3, PAR6, and aPKC), the Crumbs complex, ERM proteins (ezrin, radixin, and moesin) were recruited to establish a mature apical cap ^8–12^. In out layer polarized cells, the Hippo signaling pathway component Angiomotin (Amot) becomes apically localized. This allows YAP to translocate to the nucleus, associate with Tead4, and induce the expression of trophectoderm (TE)-specific genes, which ultimately leads to the establishment of the TE lineage ^4,13^. Interestingly, ectopic overexpression of Tfap2c, Tead4, and activated RhoA can initiate embryonic polarization in advance in 4-cell stage by regulating actin remodeling and the lateral mobility of ezrin ^14^. Besides, asymmetrically distribution of intermediate filaments, such as keratins (KRT8/18) and Lamin A/C also can promote apical polarization and TE fate determination ^15,16^. However, whether intrinsic signaling pathways are involved in the initiation of embryo polarization is less known.

During the embryo development, the fertilized egg and blastomeres till 8-cell stage have the potential to develop into both embryonic and extra-embryonic tissues. This developmental potential is known as totipotency^17^. Following the first cell fate determination, mouse epiblast cells (EPI) of the ICM retain the ability to differentiate into various tissues within the embryo but cannot give rise to extra-embryonic tissues. This restricted potential is termed pluripotency ^18^.Therefore, mESCs derived from the EPI are classified as pluripotent stem cells ^19^. Interestingly, various types of totipotent-like cells which have been demonstrated the ability to contribute into both embryonic and extra-embryonic tissues in chimeras were induced from mESCs, such as 2C-like cells (2CLC) ^20–22^, totipotent blastomere-like cells (TBLCs) ^23^, totipotent-like stem cells (TLSCs) ^24^, and chemically induced totipotent stem cells (ciTotiSCs) ^25^. Some types of these totipotent-like cells have been shown to self-organize into blastocyst-like structures, providing a valuable in vitro model system for studying pre-implantation development ^26,27^. Unexpectedly, our results showed that mESCs cultured in S/L medium can self-assemble into blastocyst-like structures when maintained in a 3D blastoid-forming medium (BFM) culture system. Further analysis revealed that CHIR99021 is essential for the formation of blastocyst-like structures, from mESCs or from totipotent-like TBLCs. CHIR99021 upregulates the expression of polarization-associated genes to promote cell polarization and TE differentiation through activation of the WNT/β-catenin signaling pathway. Additionally, Wnt/β-catenin signaling also regulates embryonic polarization *in vivo*.

## Results

### Blastocyst-like structures formed by self-assembly of mESCs

Unlike the totipotent-like cells, both our data and previous work showed that mESCs cultured in 2i/L conditions (2iL-ESCs) were unable to form blastocyst-like structures in the blastoid-forming medium (BFM) (Figure 1A and S1A) ^28,29^. Pluripotent mouse embryonic stem cells (mESCs) normally form a solid sphere structure when aggregated in serum medium, named embryoid bodies (EBs) which contain three embryonic lineages of a body upon self-differentiation. Unexpectedly, when we aggregated ESCs cultured in serum/LIF condition (SL-ESCs) in BFM for six days, around 59% cell aggregates self-assemble into the blastocyst-like structures with the fluid-filled cavities (Figure 1A and 1B). Immunostaining revealed that these structures, like natural embryos, contained SOX2-expressing cells inside of the spheres, surrounded by a peripheral layer of CDX2-expressing TE-like cells (Figure 1C and S1B). Whereas, although GATA6 positive cells were detected, most of them located at the out layer of blastoids where CDX2 expressed, with only few of them in the inner cell mass (ICM) (Figure 1C and S1B). In addition, no SOX17-positive cells were detected (Figure 1D and S1C), suggesting the primitive endoderm (PrE) lineage in the solely ESCs-derived blastoids (E-blastoids) were not well established. Time-lapse imaging revealed that CDX2 expression was activated as early as day 2 in the cell aggregates from SL-ESCs, coincident with the downregulation of NANOG (Figure S2A). In contrast, only few CDX2 positive cells were found in the aggregates at day 6 from 2iL-ESCs (Figure S2B). To further investigate the similarity in transcriptional profiles between E-blastoids and natural embryos, single-cell RNA sequencing (scRNA-seq) was performed on day 6 E-blastoids. UMAP analysis revealed that the transcriptional profiles of day 6 E-blastoids clustered closely with those of natural E3.5–E4.5 embryos (Figure 1E). Subsequent transcriptomic clustering demonstrated that the transcriptional profiles of E-blastoids co-localized with those of the TE, EPI, and PrE cells in natural embryos (Figure 1F). Analysis of lineage-specific marker genes indicated that cells in E-blastoids mainly divided into three clusters, which expressed EPI markers (*Nanog*, *Sox2*, *Pou5f1/Oct4*), TE markers (*Cdx2*, *Gata3*, *Eomes*) and PrE markers (*Gata6*, *Gata4*, *Sox17*) (Figure 1G). Yet, supporting the above staining results, a considerable proportion of PrE lineage cells also expressed *Cdx2,* suggesting their un-committed cell fate. Collectively, these findings suggest that under the BFM condition, SL-ESCs can promote the specification of the TE lineage and self-organize into blastocyst-like structures.

**Figure 1.**
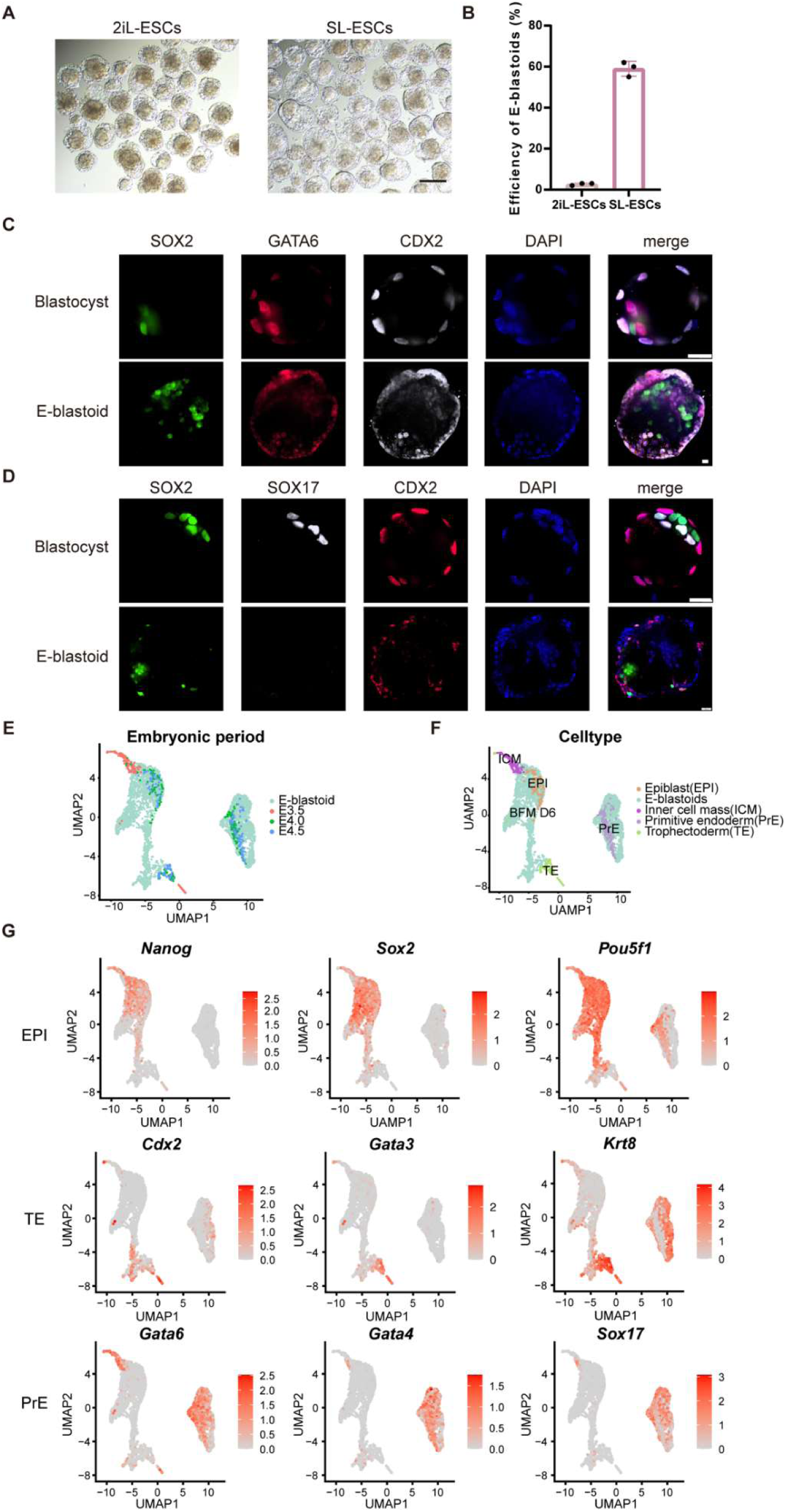
Generation of E-blastoids from mESCs. **A.** E-blastoids formed by aggregated mESC cultured in 2i/Lif (2iL) or serum/Lif (SL) medium. Scale bar: 200 μm. **B.** Quantification of the efficiency of E-blastoids formation, values are mean ± SD, each dot represents a replicate, n = 3 biological replicates in total. **C.** Representative immunofluorescence sections of E-blastoids showing the expression of EPI marker (SOX2), TE marker (CDX2), and PrE marker (GATA6). Nuclei were stained with DAPI. Scale bar: 20 μm. **D.** Representative immunofluorescence sections of E-blastoids showing the expression of EPI marker (SOX2), TE marker (CDX2), and PrE marker (SOX17). Nuclei were stained with DAPI. Scale bar: 20 μm. **E.** A UAMP plot of cells from E3.5-E4.5 natural embryos and E-blastoids. **F.** A UAMP plot showed overlap of different cell types in natural embryos and E-blastoids. **G.** Expression of EPI markers (*Nanog*, *Sox2*, and *Pou5f1*), PrE markers (*Gata6*, *Gata4*, *Sox17*) and TE markers (*Cdx2*, *Gata3* and *Eomes*) for E-blastoids and embryos scRNA-seq library.

### CHIR99021 is essential for both ESCs and totipotent-like cells to generate blastocyst-like structures

It is notable that SL-mESCs form blastocyst-like structures rather than EBs in the BFM, suggesting that chemicals (FGF4, BMP4, A83-01 and CHIR99021) in the BFM induced TE lineages in ESCs. To assess this, we aggregated SL-ESCs in the BFM, serum only or serum plus chemicals. Polarization marker aPKC, and CDX2 were detected by immunostaining at the day 3. Interestingly, while cells in serum showed no expression of aPKC and CDX2, CDX2 positive cells emerged in the aggregates in BFM or serum plus chemicals medium (Figure 2A). Moreover, aPKC polarized at the apical domain of aggregates in BFM or serum plus chemicals (Figure 2A). To further examine the role of each chemical in the BFM, individual components were omitted and their effects on the formation of blastoids were evaluated. It was observed that while removal of FGF4/heparin, BMP4, and A83-01 decrease the formation efficiency of blastoids, they did not completely abolish the ability of mESCs to form blastoids. In contrast, the removal of CHIR99021 (CHIR) led to a marked impairment in cavity formation, with the efficiency decreasing to nearly 2% (Figure 2B, C). Immunostaining of day 3 aggregates revealed that the removal of FGF4/ heparin did not significantly alter aPKC distribution or CDX2 activation (Figure 2D), consistent with the previous results that FGF4 signaling is not essential for the specification of TE lineage in pre-implantation embryos ^30^. Removal of BMP4 exhibited no apparent effect on CDX2 expression, nevertheless it attenuated the polarization of aPKC. In contrast, removal of A83-01 impaired both CDX2 activation and aPKC polarization. Strikingly, removal of CHIR99021 completely abolished CDX2 expression and aPKC polarization in nearly all aggregates, resulting in almost null blastocyst-like structure formation (Figure 2C and 2D). Moreover, supplementation of BFM basal medium with CHIR99021 alone restored detectable aPKC and CDX2 expression by day 3 and recovered blastoids formation to approximately 10% efficiency (Figure 2C and 2E). Collectively, these results establish CHIR99021 as a critical reagent regulating the polarization and the formation of E-blastoids.

**Figure 2.**
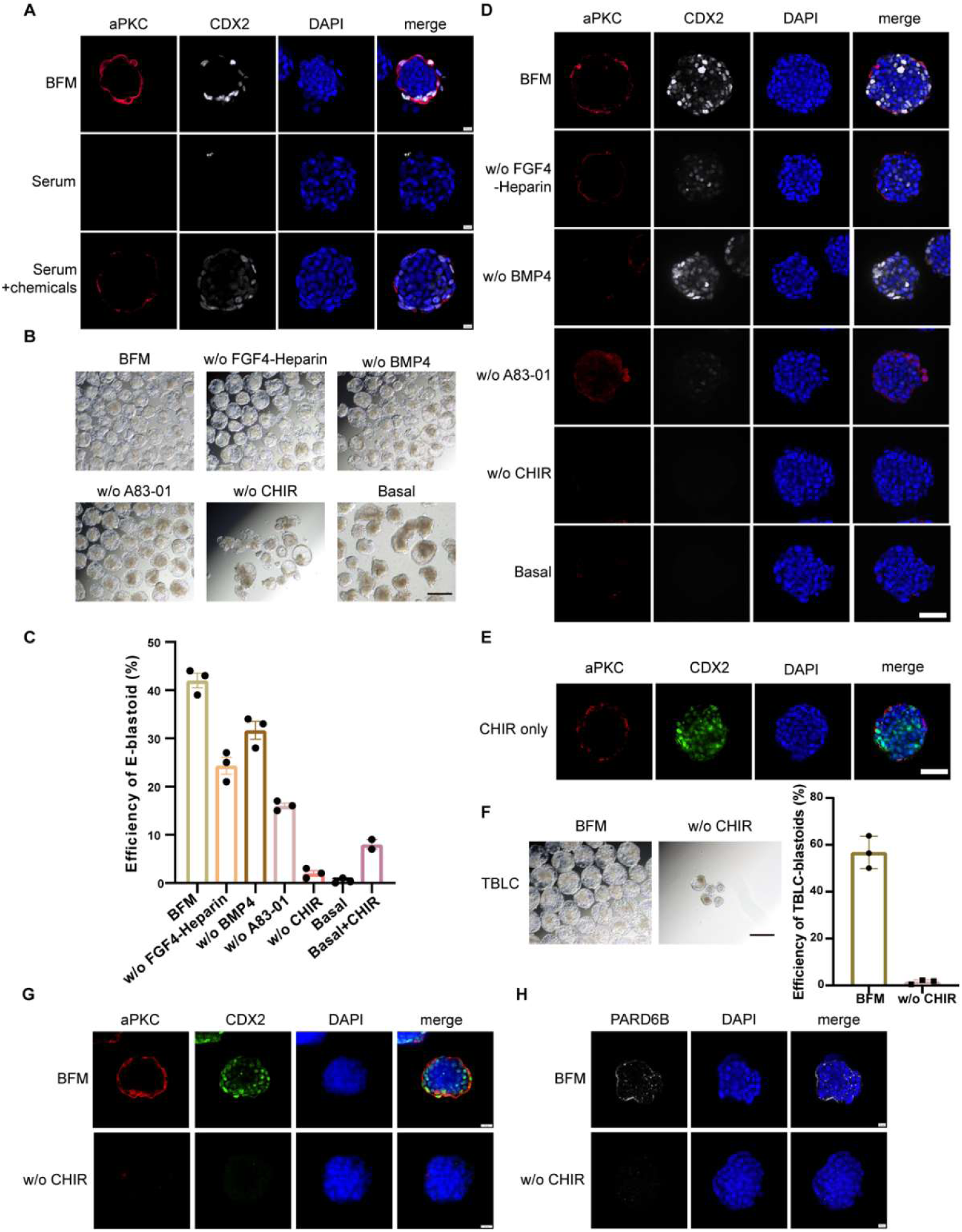
CHIR99021 is essential for the E-blastoids formation. **A.** Immunofluorescence images of cell aggregates from mESCs at day 3 showing the expression of polarity marker aPKC and TE marker CDX2. Nuclei were stained with DAPI. Scale bar: 10 μm. **B.** Microscopy of E-blastoids generated in BFM or BFM without indicated different growth factors. w/o, without. Basal, basal medium without any cytokines. Scale bar: 200 μm. **C.** Quantification of E-blastoid formation efficiency in indicated conditions. w/o, without. values are mean ± SD, n = 3 biological replicates. **D.** Immunofluorescence images of day3 aggregates showing the expression of aPKC and CDX2 in the indicated conditions. Scale bar: 50 μm. **E.** Immunofluorescence images of day3 aggregates cultured in basal medium with CHIR99021 only showing the expression of aPKC and CDX2. Scale bar: 50 μm. **F.** Microscopy of E-blastoids from TBLCs cultured in BFM or BFM without (w/o) CHIR99021 medium (Left). Scale bar: 200 μm. Quantification of TBLC-blastoids formation efficiency in indicated medium (Right); values are mean ± SD, n = 3 biological replicates. **G.** Immunofluorescence images of day 3 TBLC aggregates in indicated conditions showing the expression of aPKC and CDX2. Scale bar: 10 μm. **H.** Immunofluorescence images of day 3 TBLC aggregates in indicated conditions showing the expression of PARD6B. Scale bar: 10 μm.

To investigate whether CHIR99021 is also essential for the blastoid formation from totipotent-like cells, TBLCs ^31^ were aggregated in either BFM or BFM without CHIR99021. The results revealed that TBLCs cultured in BFM efficiently generated blastoids, consistent with previous reports ^26,27^. In contrast, withdrawal of CHIR99021 nearly completely abrogated the formation of blastocyst-like structures (Figure 2F).

Immunofluorescence staining of day 3 TBLCs aggregates revealed a complete absence of CDX2 expression, as well as a lack of apical enrichment of aPKC and PARD6B, in CHIR99021-removal condition (Figure 2G-H). These findings indicate that CHIR99021 is essential for both cellular polarization and TE marker activation in the process of blastocyst-like structure formation from mESCs as well as totipotent-like cells.

### CHIR99021 modulates the expression of TE markers and cytoskeletal dynamics during SL-mESC aggregation

To elucidate the transcriptomic effects of CHIR99021 on trophectoderm (TE) specification during blastoid formation, single-cell RNA sequencing (scRNA-seq) was performed on day 1 and day 2 aggregates cultured in either BFM or BFM without CHIR99021 (w/o CHIR) medium. At day 1, cells from the two conditions formed transcriptionally distinct clusters in the uniform manifold approximation and projection (UMAP) embedding, indicating a profound global transcriptomic divergence induced by CHIR99021 withdrawal (Figure 3A). Differential expression analysis (|log_2_FC| > 1, p < 0.05) identified 490 upregulated genes—including the TE specific factors *Cdx2*, *Gata2*, *Krt23*, and *Tfap2c*—and 329 downregulated genes in BFM-cultured cells relative to w/o CHIR counterparts (Figure 3B). KEGG pathway enrichment analysis of the 490 upregulated genes revealed significant enrichment of the Wnt signaling pathway, as well as cytoskeletal organization and Hippo signaling pathways, both of which are functionally linked to TE lineage commitment (Figure 3C). At day 2, UMAP analysis revealed that while a minor subpopulation (∼9%) of w/o CHIR cells (cluster 2) localized closed to BFM cells (cluster 0), the predominant w/o CHIR population (cluster 1) formed a transcriptionally distinct cluster clearly separated from BFM samples (Figure 3D). Differential gene expression analysis at day 2 identified 998 genes significantly upregulated in BFM samples relative to w/o CHIR conditions (|log_2_FC| > 1, p < 0.05), including the TE-associated transcription factor *Eomes* and the cytoskeletal genes *Krt8*, *Lmna* (encoding Lamin A/C), and *Myo6* (Figure 3E). In contrast, only 50 genes were downregulated. KEGG enrichment analysis demonstrated that the upregulated gene sets were also predominantly associated with cytoskeletal organization and Hippo signaling pathways (Figure 3F). Notably, among the recurrently enriched cytoskeletal regulators, KRT18 and Lamin A/C are type I and type V intermediate filament proteins, respectively, that have previously been demonstrated to be associated with TE lineage commitment ^15,16^. To validate these transcriptomic findings at the protein level, we performed immunofluorescence staining for Lamin A/C and KRT18 on day 3 aggregates cultured in BFM or w/o CHIR99021 conditions. Immunostaining revealed markedly elevated expression of Lamin A/C and KRT18 in outer cells relative to inner cells of BFM aggregates (Figure 3G). This spatially polarized expression pattern is consistent with observations in preimplantation embryos, where Lamin A/C and KRT8 protein are preferentially enriched in the outer TE layer relative to the inner cell mass (ICM) to promote CDX2 expression and TE lineage commitment during early embryogenesis ^15,16^. Notably, this outer-to-inner differential expression was markedly attenuated in w/o CHIR99021 aggregates (Figure 3G). Collectively, these data demonstrate that CHIR99021 activate TE-associated transcriptional programs and cytoskeletal organization in mESC aggregates.

**Figure 3.**
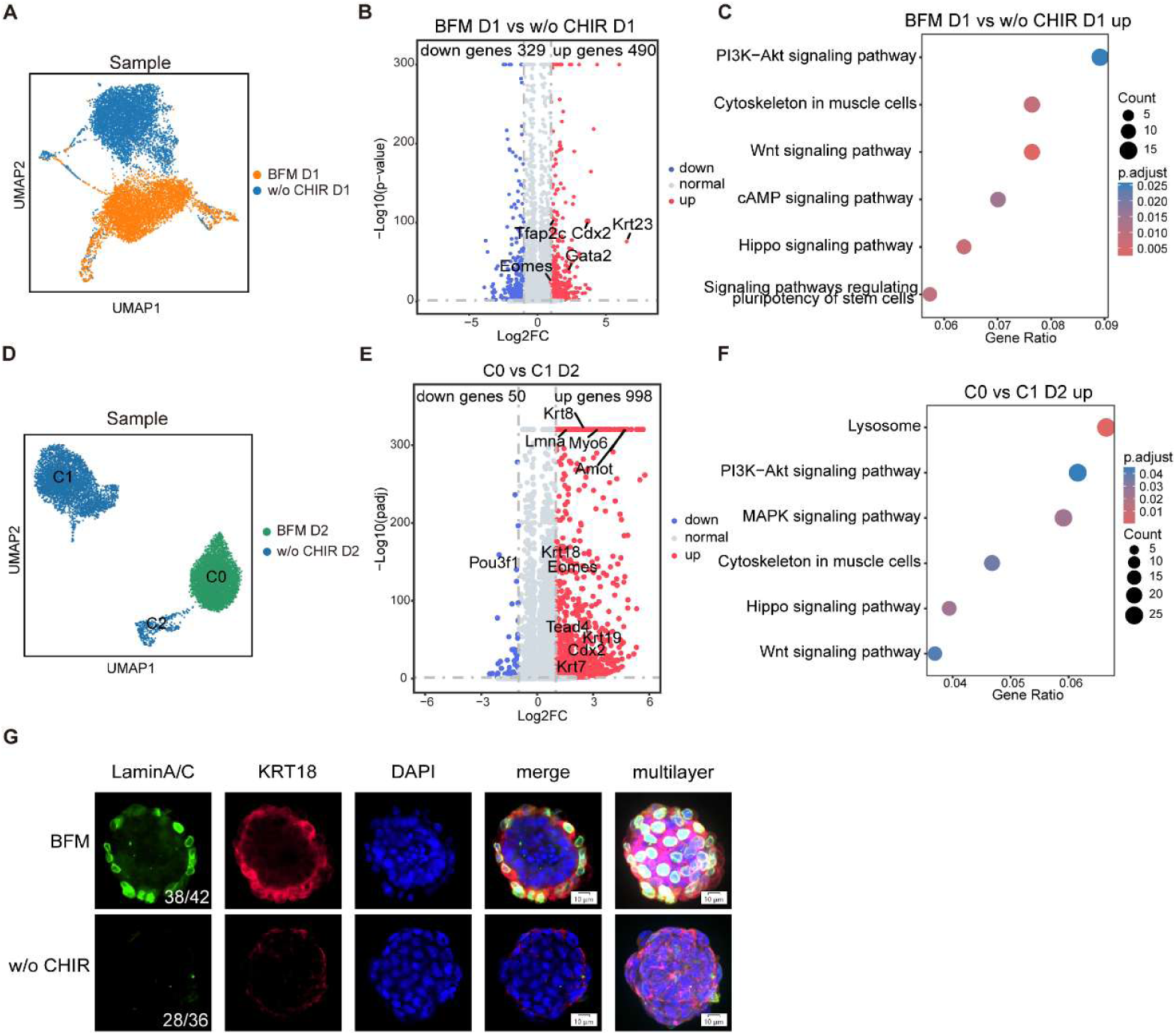
CHIR99021 modulates the expression of TE markers and cytoskeletal dynamics during mESC aggregation. **A.** UMAP visualization showing distinct cell clusters in BFM and BFM w/o CHIR99021 at day 1. **B.** Dot plots illustrating differentially expressed genes in day1 BFM and BFM w/o CHIR99021 samples. **C.** KEGG analysis of upregulated differentially expressed genes in cells in BFM at day 1 compared to those in the BFM w/o CHIR99021. **D.** UMAP visualization showing distinct cell clusters in BFM and BFM w/o CHIR99021 at day 2. **E.** Dot plots illustrating differentially expressed genes in C0 and C1 clusters in panel D. **F.** KEGG analysis of up differentially expressed genes in C0 compared to C1 clusters in panel D. **G.** Immunofluorescence images of day3 aggregates in BFM and BFM w/o CHIR99021 conditions showing the expression of cytoskeletal markers LaminA/C and KRT18. Scale bar: 10 μm.

### β-catenin regulate the formation of E-blastoids

CHIR99021, as an inhibitor targeting GSK3, can stabilize β-catenin and activate the canonical Wnt signaling pathway ^32^. To explore whether CHIR99021 functions via the β-catenin signaling pathway to induce the formation of blastoids, β-catenin levels were assessed during mESCs aggregation. Immunofluorescence staining of day 3 aggregates revealed significantly higher β-catenin levels in the BFM samples compared to the BFM without CHIR99021 samples (Figure 4A-B). We then replaced CHIR99021 in the BFM medium with Wnt3a, a ligand of the canonical Wnt pathway. After seven days of culture, the Wnt3a-treated group also generated blastocyst-like structures, although with less efficiency than that in the BFM condition (20% VS 40%) (Figure 4C). This result support that CHIR99021 functions at least partially via the canonical β-catenin signaling pathway to induce the formation of blastocyst-like structures.

**Figure 4.**
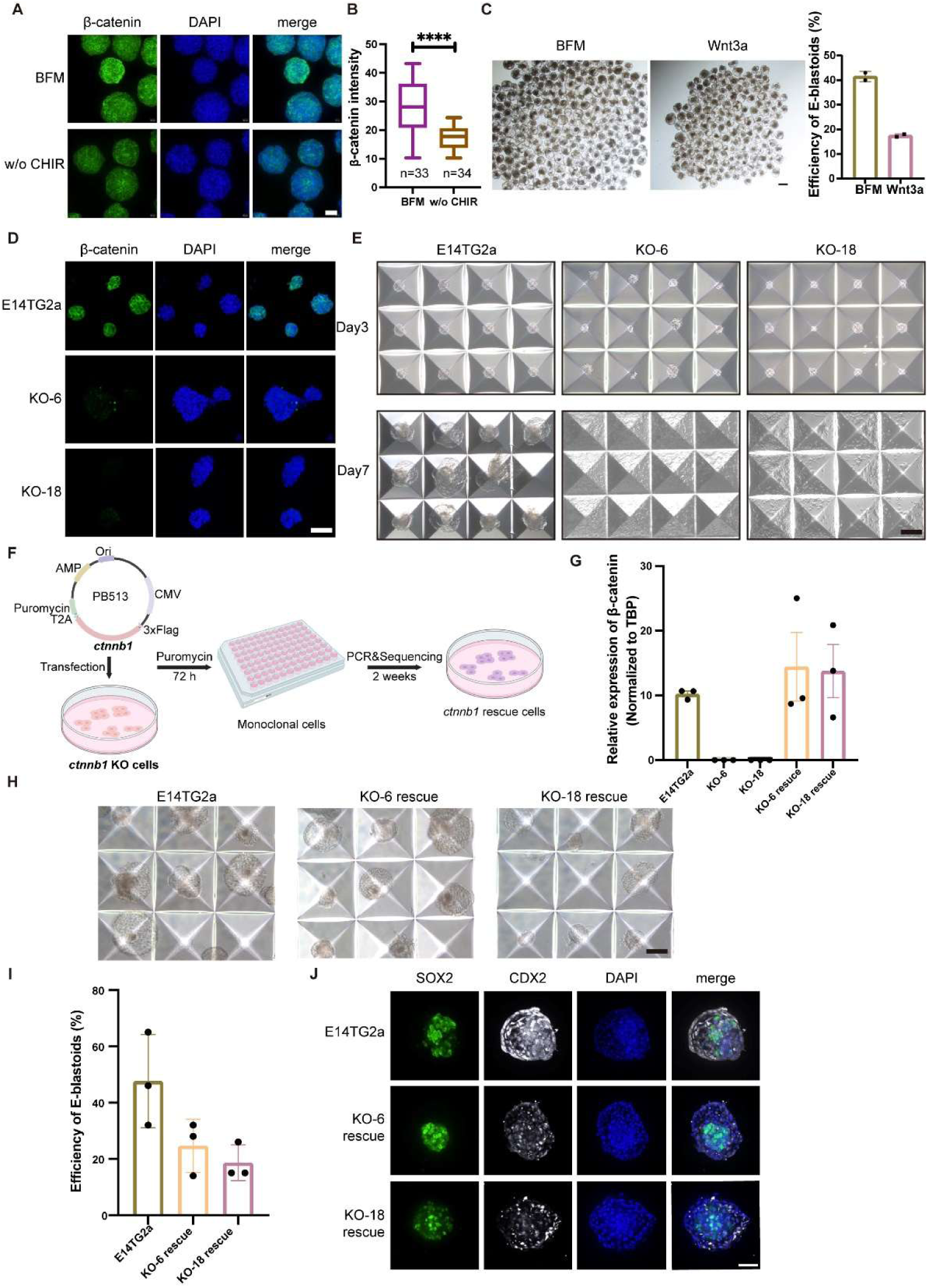
β-catenin regulates the formation of E-blastoids. **A.** Immunofluorescence images of day3 aggregates cultured in BFM and BFM w/o CHIR99021 media showing the expression of β-catenin. Nuclei were stained with DAPI. Scale bar: 50 μm. **B.** Quantification of β-catenin fluorescence intensity in the indicated conditions; n number is indicated. Values are mean ± SD, unpaired t-test, **** p < 0.0001 **C.** The formation of E-blastoids in BFM or BFM medium with CHIR99021 replaced by Wnt3a (Wnt3a) (Left). Scale bar: 200 μm. The bar chart illustrates the efficiency of E-blastoid formation in the indicated medium (Right). Values are mean ± SD, n = 2 biological replicates. **D.** Immunofluorescence images of KO-6 and KO-18 clones showing β-catenin expression. Nuclei were stained with DAPI. Scale bar: 50 μm. **E.** Aggregates of wild-type (E14TG2a) and β-catenin KO cells at the indicated days. Scale bar: 200 μm. **F.** Schematic overview of establishment of Flag-β-catenin rescued cells in β-catenin KO cells. **G.** Quantitative mRNA analysis of β-catenin expression levels in KO and rescued cells; values are mean ± SE, n = 3 biological replicates. **H.** Formation of E-blastoids from wild-type (E14TG2a) and β-catenin rescued cells. Scale bar: 200 μm. **I.** Efficiency of E-blastoids formation from wild-type (E14TG2a) and β-catenin rescued cells; values are mean ± SD, n = 3 biological replicates. **J.** Immunofluorescence images of E-blastoids formed from wild-type (E14TG2a) and β-catenin rescued cells. Scale bar: 50 μm.

To further investigate the function of β-catenin in the generation of blastoids, we knocked out β-catenin in mESCs and selected two homozygous knockout clones (KO-6 and KO-18), in which frameshift mutations were generated through targeted deletion of exons 1–8 of the β-catenin gene loci (Fig. S3A-B). Immunostaining confirmed that β-catenin was undetectable in both the KO-6 and KO-18 clones (Figure 4D). β-catenin KO did not affect mESCs morphology, and proliferation (Fig. S3C), a finding consistent with previously published results ^33^. Subsequently, blastocyst-like structures formation assays were performed using β-catenin KO cells. It showed that β-catenin KO cells formed comparable aggregates at day3 compared to control wild-type cells. However, at day7, while control cells predominantly formed blastocyst-like structures, β-catenin-KO cells remained dispersed in the microwell plates and failed to organize into spherical structures (Figure 4E). These results indicate that β-catenin is essential for blastoid formation. To further validate the role of β-catenin in the formation of blastocyst-like structures, Flag-β-catenin was reintroduced into β-catenin KO cells (Figure 4F). QPCR analysis was conducted to evaluate β-catenin mRNA expression levels. Two clones with

β-catenin expression levels comparable to wild-type (WT) mESCs were selected for further analysis (Figure 4G). Immunofluorescence staining with both anti-FLAG and anti-β-catenin antibodies collectively confirmed successful reconstitution of β-catenin in rescued cells (Figure S3D). Subsequently, aggregation assays were conducted using wild-type and rescued cells. It was observed that β-catenin ectopically expressing cells restored blastocyst-like structure generation with an efficiency of ∼20% under BFM condition (Figure 4H and I). Immunostaining analysis revealed that the blastocyst-like structures derived from β-catenin rescue cells exhibited morphological features comparable to those of blastoids from wild-type ESCs, with inner SOX2-positive epiblast-like cells and a cavity surrounded by CDX2-positive TE-like cells (Figure 4J). These results provide conclusive evidence that β-catenin functions as a critical signaling molecule in the formation of E-blastoids.

### β-catenin regulates the cell polarization of blastocyst-like structures

To investigate the impact of β-catenin on cell polarization during E-blastoids formation, immunostaining for polarity markers was conducted on day 3 aggregates derived from β-catenin KO and rescued cells. In controls, PARD6B localized to the apical regions of outer cells, consistent with previous studies indicating that E-blastoid recapitulate the polarization patterns observed in natural embryos. In contrast, β-catenin KO aggregates showed no β-catenin expression or apical enrichment of PARD6B, suggesting that polarization does not occur during aggregation in the absence of β-catenin (Figure 5A). Importantly, restoration of β-catenin in KO cells rescued apical localization of PARD6B comparable to that seen in the control group (Figure 5A). These findings collectively demonstrate that β-catenin is critical for establishing cellular polarization during E-blastoids formation.

**Figure 5.**
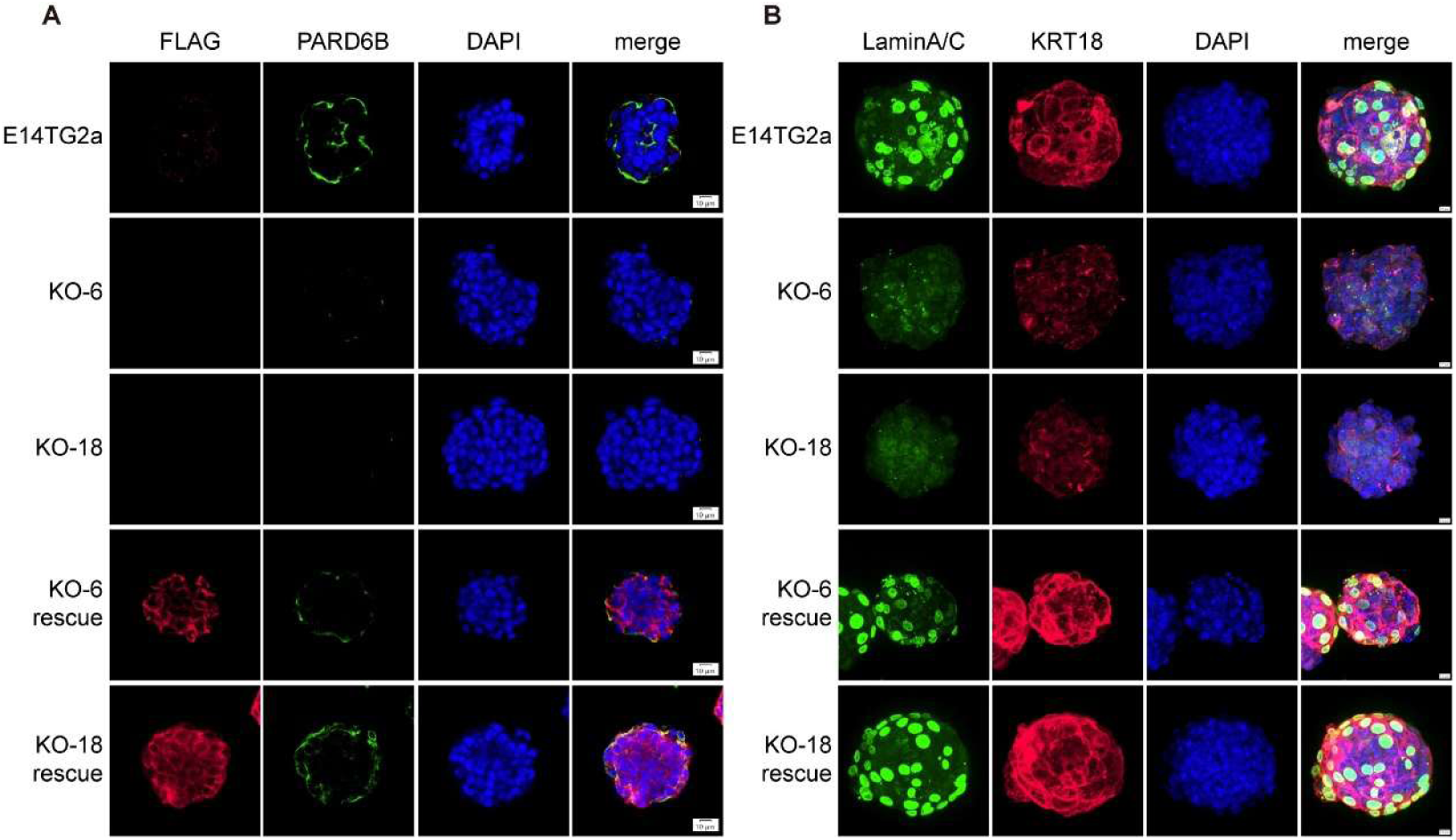
β-catenin regulate the polarization of aggregates. **A.** Immunofluorescence staining of FLAG and PARD6B at day 3 during the formation of E-blastoids from wild-type (E14TG2a), β-catenin KO and rescued cells. Scale bar: 10 μm. **B.** Immunofluorescence staining of LaminA/C and KRT18 at day 3 during the formation of E-blastoids from wild-type (E14TG2a), β-catenin KO and rescued cells. Scale bar: 10 μm.

LaminA/C and KRT18 expression were also analyzed in day 3 aggregates derived from β-catenin KO and rescued cells. In wild-type controls, the expression and distribution of Lamin A/C and KRT18 were consistent with prior observations (Figure 3G). In contrast, in two independent β-catenin KO clones, Lamin A/C was uniformly distributed across the nuclei of all cells within the aggregates, with no apparent spatial bias in expression. KRT18 expression was markedly diminished and evenly dispersed throughout the KO aggregates, lacking any discernible inner–outer polarity. In contrast, restoration of β-catenin rescued periphery expression of Lamin A/C and KRT18 (Figure 5B), indicating that β-catenin regulates the expression and remodeling localization of these proteins, thereby influencing TE fate specification in mESCs aggregates.

### β-catenin regulate the polarization *in vivo*

*In vivo*, β-catenin is a maternal factor, expressed across from oocytes to blastocyst stage^34^. Previous work revealed that although maternal-zygotic deletion of β-catenin (exons 8-13) reduces blastocyst numbers at E4.5 below the Mendelian ratio, it does not prevent blastocyst formation. Nevertheless, it is essential for implantation and subsequent development, concurrent with impaired CDX2 expression ^34–36^. Albeit with these results, the function of β-catenin in embryo polarization is less known. To assess the role of β-catenin in early embryo development, we employed the Trim-away method to degrade β-catenin protein in embryos ^37^(Figure 6A). Staining of β-catenin at 2-Cell stage confirmed that β-catenin was abolished in the embryos injected with both antibodies and Trim21 mRNAs but not in the control embryos which injected with only Trim21 mRNAs (Figure 6B). Following the depletion of β-catenin, no significant effect was observed on the 2-cell developmental rate of embryos; however, a reduction of blastocyst formation rate was noted (Figure 6C and S4A). Moreover, we noticed the less compact of the β-catenin-Trimaway embryos attributed to the disturbance of adherent junctions ^38^. qPCR analysis performed of the 8C embryos demonstrated that reduction in β-catenin protein levels effectively suppresses transcription of the *Pard6B*, *Krt8*, *Amot*, and *Lmna* genes (Figure 6D). Immunofluorescence staining demonstrated that the absence of β-catenin led to a significant decrease in the expression level of aPKC and also disrupted the organization of F-actin (Figure 6E-F). The expression levels of KRT18 and Lamin A/C were also downregulated in the Trim-away samples (Figure 6G and S4B). To assess the impact of impaired cell polarities on TE fate, we analyzed the expression of the TE markers CDX2 and YAP. CDX2 and nuclear YAP expression was markedly reduced in β-catenin depletion samples. (Figure 6H, S4C and S4D). These results suggested that β-catenin elimination in early embryos downregulated the cell polarity-related genes and therefore impaired the TE-lineage specification.

**Figure 6.**
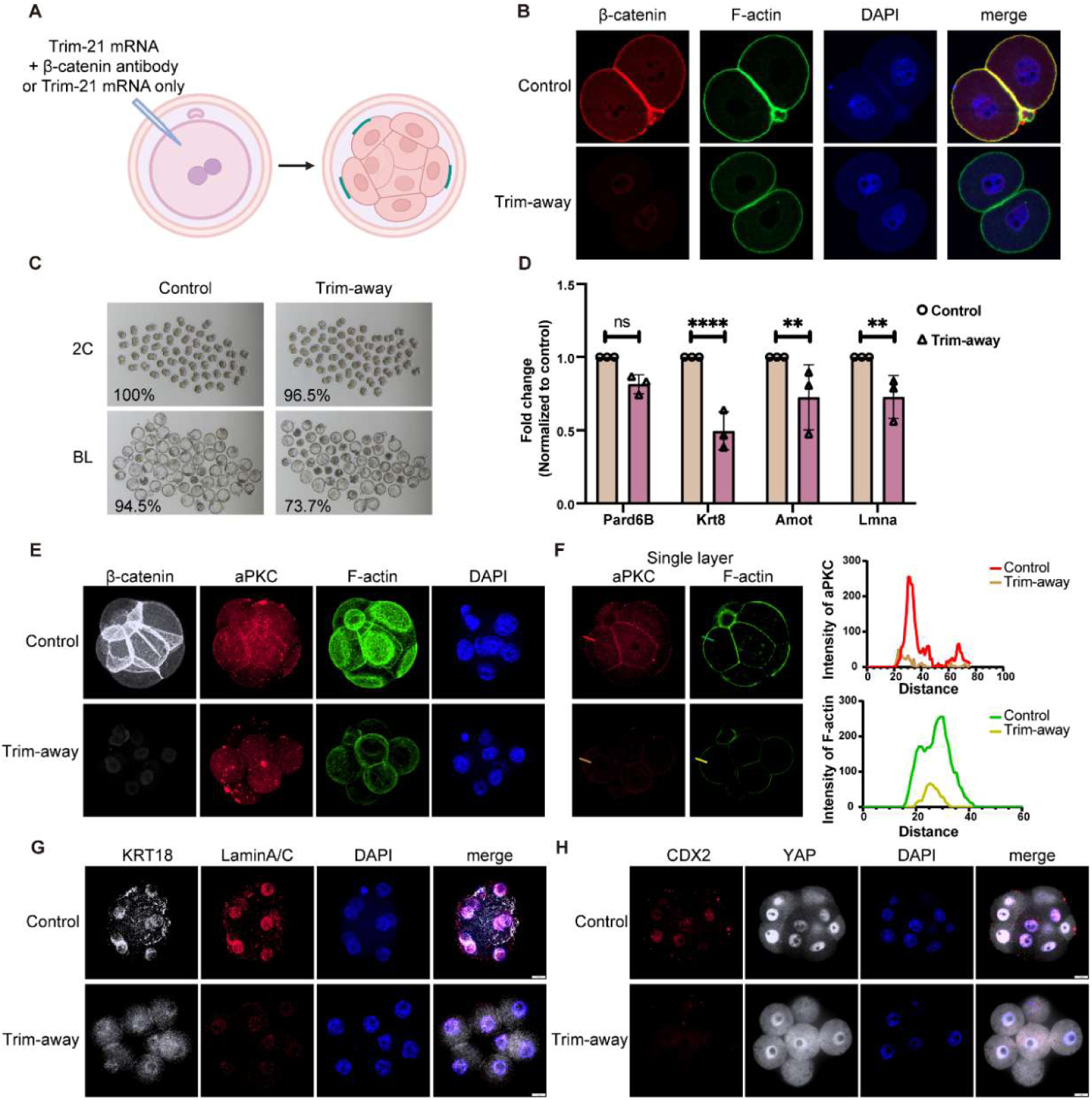
β-catenin regulates embryonic polarization *in vivo*. **A.** Schematic overview of β-catenin Trim-away in zygotes. **B.** Immunofluorescence staining of aPKC and F-actin in control and Trim-away embryos at 2-Cell stage. **C.** Representative microscopies of embryos at 2-Cell and blastocyst stage from the control and Trim-away group. **D.** Quantitative mRNA analysis of indicated gene expressions in control and Trim-away samples; values are means ± SD, paired t-test, ns, p = 0.050536, ****, p = 0.00003, **, p = 0.006505 (Amot), **, p = 0.006764 (Lmna), n = 3 biological replicates. **E.** Immunofluorescence staining of aPKC and F-actin in control and Trim-away embryos at 8C to morula l stage. Scale bar: 10 μm. **F.** Fluorescence intensity profile of aPKC and F-actin shown along the white line in control and Trim-away embryos at 8C to morula stage (showed in left). **G.** Immunofluorescence staining of KRT18 and LaminA/C in control and Trim-away embryos at 8C to morula stage. Scale bar: 10 μm. **H.** Immunofluorescence staining of CDX2 and YAP in control and Trim-away embryos at 8C to morula stage. Scale bar: 10 μm.

## Discussion

Owing to the limit of embryonic materials, research on *in vitro* synthetic embryo models has been a steady increase in recent years. Mouse ESCs, a type of pluripotent stem cells, can differentiate into various cell types that constitute the embryo. However, they are unable to autonomously differentiate into extraembryonic cell types. Unexpectedly, our experiments revealed that mESCs cultured in SL medium can self-assemble into blastocyst-like structures when maintained in a 3D BFM culture system. These structures exhibit some key features of natural embryos, including the embryo polarization and the initiation of first cell fate determination, which leads to the formation of TE-like cells. Further analysis revealed that CHIR99021 in BFM is essential for the formation of blastocyst-like structures from both mESCs and totipotent-like TBLCs aggregates. CHIR99021, as an inhibitor of GSK3α/β, upregulate intermediate filament proteins and cytoskeleton-remodeling genes to promote cell polarization and TE differentiation via canonical Wnt/β-catenin pathway. Importantly, β-catenin also regulate the embryo polarization *in vivo*. Embryos without β-catenin impaired the level of polarization-related proteins and the development of blastocyst. However, except its role in regulating downstream gene expression by interacting with transcription factors, β-catenin also function together with α-catenin to stabilize the adherent junctions ^39,40^. Nevertheless, previous studies have shown that E-cadherin knockout does not impair embryonic polarization and CDX2 expression ^41^, suggesting that β-catenin regulates polarity proteins likely through the Wnt/β-catenin signaling pathway rather than the impairment of adherent junctions. However, whether these effects stem from direct transcriptional control of polarity determinants or secondary signaling cascades requires further investigation.

Interestingly, previous study revealed that CHIR99021 is critical for efficient generation of blastocyst-like structure from the aggregation of ESCs and TSCs ^42^. CHIR99021 or LY2090314, another Wnt signaling activator, were also a major component to induce cells expressed bi-potential markers from ESCs in recent studies ^43,44^. These results support the critical role of β-catenin in the activation of TE lineage and the formation of blastoids. Furthermore, our result showed CHIR99021 is indispensable for the formation of blastoids from totipotent-like TBLC cells. Our result suggest that we should be more prudent to use the blastoid generation as a functional assay for evaluating the totipotent-like cells. Meanwhile, although totipotent-like cells exhibit certain features reminiscent of natural embryos, unlike, natural embryos, extrinsic signaling is essential for their ability to generate blastocyst-like structures, implying that additional extracellular signaling cues may be required to achieve fully functional blastoid development.

## Methos and Materials

### Mice

Animal experiments were performed in accordance with the Guide of the Care and Use of Laboratory Animals by the National Research Council and approved by Institutional Animal Care and Use Committee of Guangzhou National Laboratory. Mice were purchased from ZHUHAI BESTEST BIOTECHNOLOGY CO., LTD. and housed in environmentally controlled room.

### Cell lines

Mouse embryonic stem cell line, E14Tg2a was employed for this study. TBLCs, β-catenin KO cell lines and rescued cell lines were generated from E14tg2a. Cells were cultured in Dulbecco’s Modified Eagle Medium (DMEM, high glucose) supplemented with 15% fetal bovine serum (FBS), 1× non-essential amino acids (NEAA), 1× GlutaMAX, 0.1 mM β-mercaptoethanol, and 10³ U mL⁻¹ recombinant human leukemia inhibitory factor (LIF). Cells were maintained at 37°C in an atmosphere incubator containing 5% CO₂. mESCs were seeded onto 0.1% gelatin-coated plates and passaged every two days using 0.05% trypsin-EDTA at a 1:10 splitting ratio.

### Generation of E-blastoid

The methodology for generating E-blastoid formation is consistent with previously reported protocols ^29^. AggreWell^TM^400 plates (STEMCELL Technologies, 34415) were prepared according to the manufacturer’s instructions. MESCs colonies were dissociated into single cells using 0.1% trypsin-EDTA for 5 minutes at 37 °C. TBLCs were resuspended following dissociation and transferred to plates coated with 0.1% gelatin, and incubated at 37 °C for at least 30 minutes to remove co-cultured MEF cells. mESCs or TBLCs remaining in suspension were collected and counted. A total of 1.8 × 10^4^ cells were seeded into each well of a 24-well AggreWell plate in BFM medium supplemented with 2 μM Y-27632. Twenty-four hours later, the medium was replaced with fresh BFM medium devoid of Y-27632, and subsequently changed every two days. The BFM medium comprised 25% TSC basal medium, 25% N2B27 basal medium, and 50% M16 medium, supplemented with 12.5 ng/mL rhFGF4, 0.5 μg/mL heparin, 5 ng/mL BMP4, 3 μM CHIR99021, and 0.5 μM A83-01. The TSC basal medium was prepared as RPMI 1640 supplemented with 20% FBS, 1% sodium pyruvate, and 0.1 mM β-mercaptoethanol. The N2B27 basal medium was prepared as a 1:1 mixture of DMEM/F-12 and Neurobasal medium, supplemented with 0.5% N-2 Supplement (100×), 0.5% B-27™ Supplement (50×), 1% GlutaMAX, 1% NEAA, and 0.1 mM β-mercaptoethanol.

### TBLC induction

The methodology used for establishing TBLCs was in accordance with previously published protocols ^31^. The TBLC medium consisted of mESCs culture medium supplemented with 2.5 nM of PlaB. To generate TBLCs, mESCs were seeded onto a feeder layer of irradiated mouse embryonic fibroblasts (MEFs) and maintained in TBLC medium. Cells were passaged every 3 to 4 days using 0.1% trypsin-EDTA at a splitting ratio of 1:3 and seeded onto MEFs. A minimum of six passages was conducted prior to the initiation of aggregation. Cells from the 8th to the 11th passage were used for the aggregation experiments.

### Cell transfection

mESCs were seeded in 0.1% gelatin-coated 6-well plates at a cell density of 6 × 10⁵ cells per well. Transfections were carried out using Lipofectamine® 3000 according to the manufacturer’s instructions. Briefly, 1 μg of plasmid DNA and 2.5 μL of P3000 reagent were diluted in 100 μL of Opti-MEM medium. In a separate tube, 2.5 μL of Lipofectamine® 3000 was diluted in another 100 μL of Opti-MEM. The diluted DNA solution was then combined with the diluted Lipofectamine® 3000 reagent and incubated for 15 minutes at room temperature to form DNA-lipid complexes. These complexes were subsequently added to the mESC cultures. 24 hours post-transfection, cells were selected with 1 μg/mL puromycin for 72 hours. For the generation of β-catenin KO cell lines, FACS-based sorting was performed following selection, and individual clones derived from single cells were isolated and analyzed by PCR and sequencing.

### CRISPR knockout of β-catenin in mESCs

β-catenin KO mESCs lines were generated by CRISPR knockout technique. SgRNAs (Aulicino et al., 2020) were annealed and subsequently cloned into the px459-SpCas9-Puro vector. Briefly, mESCs were transfected with px459-SpCas9-Puro vectors (a kind gift from Dr Ian Chambers lab) with two guide RNAs targeting the first and eighth exon of β-catenin. After transfection and drug selection, the mESCs were seeded as single clones, which were then picked up and expanded. For genotyping, genomic DNA was extracted from β-catenin KO mESCs using the Mouse Direct PCR Kit following the manufacturer’s instructions. PCR amplification was then carried out using 2× Rapid Taq Master Mix and genotype-specific primers, after which the PCR products were analyzed by gel electrophoresis. For the KO cell lines, clones that were confirmed as successfully KO through gel electrophoresis were further validated by Sanger sequencing. Homozygous KO clones exhibiting frameshift mutations were selected for subsequent culture and experiments. The gRNAs and primers used are listed in Table S1.

### Rescue of β-catenin in knockout cells

For the β-catenin rescue experiment, the β-catenin coding DNA sequence (CDS) was amplified from mESCs cDNA using PrimeSTAR Max DNA Polymerase. FLAG-tagged β-catenin CDS was generated by PCR using the wild-type product as a template and subsequently cloned into a PB513-PiggyBac-Puro-based vector using Gibson assembly follow the manufacturer’s instructions. The primers used are listed in Table S1.

Following 24 hours of transfection with the PB513-FLAG-β-catenin plasmid into KO cells, cells were exposed to 1 μg/mL puromycin selection for 72 hours, after which monoclonal isolation was carried out via flow cytometry. Total RNA was extracted from each isolated clone and analyzed by qPCR. The clone displaying β-catenin expression levels most comparable to those observed in mESCs was selected for subsequent experiments.

### Mouse embryo collection

Six- to eight-week-old C57BL/6J female mice were superovulated via intraperitoneal injection of 7.5 U pregnant mare serum gonadotropin (PMSG), followed 48 hours later by an intraperitoneal injection of 7.5 U human chorionic gonadotropin (hCG). The treated females were then bred with C57BL/6J male mice. Eighteen to twenty hours after hCG injection, the oviducts of mated females were surgically punctured to collect oocyte-cumulus complexes. Following digestion with 1 mg/mL hyaluronidase for 10–15 seconds at 37 °C, the denuded oocytes were retrieved and cultured in drops of EmbryoMax Advanced KSOM embryo culture medium under a layer of mineral oil.

### Immunofluorescence staining

E-blastoids, TBLC-blastoids, and mouse embryos were fixed with 4% paraformaldehyde (PFA) for 20 minutes at room temperature. Following fixation, samples were washed three times (5 minutes per wash) in 0.1% PBST (PBS containing 0.1% Triton X-100), permeabilized with 0.3% Triton X-100 in PBS for 30 minutes, and subsequently blocked with blocking buffer (3% donkey serum, 1% BSA, 0.1% Triton X-100 in PBS) for 2 hours at room temperature. The samples were then incubated overnight with primary antibodies diluted in blocking buffer at 4 °C. Afterward, they were washed four times (5 minutes each) with 0.1% PBST, followed by a 2-hour incubation with secondary antibodies in blocking buffer at room temperature. A final washing step was performed with 0.1% PBST four times. Nuclei were stained with DAPI, and filamentous actin was labeled with Phalloidin 488. Image acquisition was performed using Olympus Ixplore SpinSR. The antibodies used are listed in Table S2.

Coverslips with cells were fixed with 4% PFA for 20 min at room temperature, then washed with PBS 3 times (5 minutes each). The samples were blocked and permeabilized in PBS with 0.3% Triton and 1% normal donkey serum for 1 hour at room temperature. The coverslips were overnight incubated with primary antibodies in PBS at 4 ℃, washed three times with PBS and incubated with secondary antibodies for 1 hour at room temperature. Finally, coverslips were sealed on slides using Fluoromount-G medium. Image acquisition was performed using Olympus Ixplore SpinSR. The antibodies used are listed in Table S2.

### RNA extraction

Total RNA of β-catenin KO or rescue mESCs was extracted by TRIzol™ Reagent. For 6-well plate cells, added 200 μL TRIzol™ Reagent to lyse the cells, transferred cells into a clean tube. Incubated for 5 minutes then added 50 μL chloroform into the tube and incubated for 2-3 minutes, then centrifuged for 15 minutes at 12, 000× g at 4 ℃. Transferred the upper layer aqueous phase to a new tube, then added 100 μL isopropanol, incubated for 10 minutes at room temperature and centrifuged for 10 minutes at 12, 000× g at 4 ℃. The supernatant was discarded and 500 μL 75% ethanol was added to resuspend the precipitates, then centrifuged for 5 minutes at 8000× g at 4 ℃. After that, discarded the supernatant and air dried the RNA for 5-10 minutes, resuspended the RNA in 50 μL RNase-free water.

### qRT-PCR analysis

Total RNA was reverse transcribled with PrimeScript RT Reagent Kit (Takara, RR047A). qPCR with TB Green Premix Ex TaqTM II (Takara) was performed on QuantStudio Q7 system (Invitrogen) following the manufacturer’s instructions. Gene expression was normalized to TBP. The primers used are listed in Table S1.

### Trim21 mRNA preparation

The pGEMHE-mCherry-mTrim21 plasmid was linearized using PacI restriction enzyme, followed by the addition of 0.5 M EDTA and 3 M sodium acetate to precipitate the linearized DNA. After quantification, 1 μg of the purified plasmid was used for in vitro transcription with the mMESSAGE mMACHINE® T7 Ultra Kit according to the manufacturer’s instructions. The synthesized mRNA was precipitated with LiCl, and its concentration was determined spectrophotometrically. Finally, the mRNA was aliquoted into tubes at 6 μL per aliquot and stored at –80°C.

### Embryo manipulation

Female B6D2F1 mice aged eight to nine weeks were induced to undergo superovulation through the administration of 10 U PMSG, followed by an intraperitoneal injection of 14 U hCG 48 hours later. Following hCG administration, the superovulated females were paired with adult male B6D2F1 mice for mating. Zygotes were collected from the oviducts and cultured in KSOM medium Millipore at 37°C under 5% CO2 in a humidified incubator. Purified Trim21 mRNA was mixed with β-catenin antibody to achieve final concentrations of 400 ng/μL for Trim21 mRNA and 0.125 mg/mL for the β-catenin antibody. The mixture was microinjected into the cytoplasm of fertilized eggs using an Eppendorf FemtoJet 4i. Embryos at the late 8-cell stage were harvested 72–74 hours post-hCG injection and prepared for subsequent analyses.

### Single-cell RNA-seq data processing

Raw sequencing data based on the 10x Genomics library method were first processed using TrimGalore (v0.6.5) to remove adapter sequences and low-quality bases. Then cleaned reads were mapped to the mouse reference genome (mm10) using STAR (v2.7.6a). The cells with a minimum cut-off of 2000 genes or over 10% proportion of UMI counts from mitochondrial genes were removed. Cells with total UMI counts above 100000 were also filtered. The python package scrublet (v0.2.3) was used to remove potential cell doublets. Briefly, normalization and scaling were performed on the raw count matrix. Non-linear dimension reduction using uniform manifold approximation and projection (UMAP) was subsequently performed with the top 30 principal components (PCs) as input. Differential expressed genes were calculated using Seurat packages (v5.1.0) FindMarkers function (log2FC >1, adjust p value < 0.05).

The integrated analysis with embryo samples were using the canonical correlation analysis (CCA) pipeline in Seurat to remove batch effects. Scale data with 2,000 highly variable genes were used to compute PCA, and 30 principal components were employed to calculate the UMAP coordinates with RunUMAP. We performed Kyoto Encyclopedia of Genes and Genomes (KEGG) enrichment analysis using R package clusterProfiler (v4.10.0).

### Statistical analysis

Quantification and statistical analysis were performed using GraphPad Prism 9.0 software. Data are presented as the mean ± SD or mean ± SE. The p values were calculated by Student’s two-sided unpaired t-test and indicated in the figures. The p values < 0.05 were considered statistically significant.

## Acknowledgments

We thank members of the Zhang laboratory for inspiring discussions and comments; the Animal Center and the flow cytometry Core of Public Technology Service Center at Guangzhou National Laboratory for their support. This work was supported by grants to M.Z. from the National Natural Science Foundation of China (grant no. 32570958), and Major Project of Guangzhou National Laboratory (grant no. GZNL2023A02005).

## Author contributions

W.C., and M.Z. conceived, designed and conducted the studies; J.H., Y.Z., B.H., S.S., J.F., W.S., L.L., and D.L., performed experiments; X.Z. performed bioinformatics analysis; J.H. and M.Z. wrote the paper, with input from all authors.

## Lead contact

Further queries and reagent requests may be directed and will be fulfilled by the lead contact, Man Zhang.

## Materials availability

All unique/stable reagents generated in this study are available from the Lead Contact with a completed Materials Transfer Agreement.

## Data and code availability

ScRNA-seq data can be accessed with the CNCB accession number: PRJCA062787.

## References

1. Chazaud, C., and Yamanaka, Y. (2016). Lineage specification in the mouse preimplantation embryo. Development 143, 1063–1074. 10.1242/dev.128314.

2. Yao, C., Zhang, W., and Shuai, L. (2019). The first cell fate decision in pre-implantation mouse embryos. Cell Regeneration 8, 51–57. 10.1016/j.cr.2019.10.001.

3. Korotkevich, E., Niwayama, R., Courtois, A., Friese, S., Berger, N., Buchholz, F., and Hiiragi, T. (2017). The Apical Domain Is Required and Sufficient for the First Lineage Segregation in the Mouse Embryo. Developmental Cell 40, 235–247.e237. 10.1016/j.devcel.2017.01.006.

4. Hirate, Y., Hirahara, S., Inoue, K.-i., Suzuki, A., Alarcon, Vernadeth B., Akimoto, K., Hirai, T., Hara, T., Adachi, M., Chida, K., et al. (2013). Polarity-Dependent Distribution of Angiomotin Localizes Hippo Signaling in Preimplantation Embryos. Current Biology 23, 1181–1194. 10.1016/j.cub.2013.05.014.

5. Buckley, C.E., and St Johnston, D. (2022). Apical–basal polarity and the control of epithelial form and function. Nature Reviews Molecular Cell Biology 23, 559–577. 10.1038/s41580-022-00465-y.

6. Zhu, M., Leung, C.Y., Shahbazi, M.N., and Zernicka-Goetz, M. (2017). Actomyosin polarisation through PLC-PKC triggers symmetry breaking of the mouse embryo. Nature Communications 8. 10.1038/s41467-017-00977-8.

7. Zhu, M., and Zernicka-Goetz, M. (2020). Building an apical domain in the early mouse embryo: Lessons, challenges and perspectives. Current Opinion in Cell Biology 62, 144–149. 10.1016/j.ceb.2019.11.005.

8. Louvet, S., Aghion, J., Santa-Maria, A., Mangeat, P., and Maro, B. (1996). Ezrin Becomes Restricted to Outer Cells Following Asymmetrical Division in the Preimplantation Mouse Embryo. Developmental Biology 177, 568–579. 10.1006/dbio.1996.0186.

9. Vinot, S., Le, T., Ohno, S., Pawson, T., Maro, B., and Louvet-Vallée, S. (2005). Asymmetric distribution of PAR proteins in the mouse embryo begins at the 8-cell stage during compaction. Developmental Biology 282, 307–319. 10.1016/j.ydbio.2005.03.001.

10. Hirate, Y., Hirahara, S., Inoue, K.i., Kiyonari, H., Niwa, H., and Sasaki, H. (2015). Par-aPKC-dependent and -independent mechanisms cooperatively control cell polarity, Hippo signaling, and cell positioning in 16-cell stage mouse embryos. Development, Growth & Differentiation 57, 544–556. 10.1111/dgd.12235.

11. Ajduk, A., and Zernicka-Goetz, M. (2016). Polarity and cell division orientation in the cleavage embryo: from worm to human. Molecular Human Reproduction 22, 691–703. 10.1093/molehr/gav068.

12. Lamba, A., and Zernicka-Goetz, M. (2023). The role of polarization and early heterogeneities in the mammalian first cell fate decision. In Cell Polarity in Development and Disease, pp. 169–196. 10.1016/bs.ctdb.2023.02.006.

13. Hirate, Y., and Sasaki, H. (2014). The role of angiomotin phosphorylation in the Hippo pathway during preimplantation mouse development. Tissue Barriers 2. 10.4161/tisb.28127.

14. Zhu, M., Cornwall-Scoones, J., Wang, P., Handford, C.E., Na, J., Thomson, M., and Zernicka-Goetz, M. (2020). Developmental clock and mechanism of de novo polarization of the mouse embryo. Science 370. 10.1126/science.abd2703.

15. Lim, H.Y.G., Alvarez, Y.D., Gasnier, M., Wang, Y., Tetlak, P., Bissiere, S., Wang, H., Biro, M., and Plachta, N. (2020). Keratins are asymmetrically inherited fate determinants in the mammalian embryo. Nature 585, 404–409. 10.1038/s41586-020-2647-4.

16. Skory, R.M., Moverley, A.A., Ardestani, G., Alvarez, Y., Domingo-Muelas, A., Pomp, O., Hernandez, B., Tetlak, P., Bissiere, S., Stern, C.D., et al. (2023). The nuclear lamina couples mechanical forces to cell fate in the preimplantation embryo via actin organization. Nature Communications 14. 10.1038/s41467-023-38770-5.

17. Condic, M.L. (2014). Totipotency: What It Is and What It Is Not. Stem Cells and Development 23, 796–812. 10.1089/scd.2013.0364.

18. Riveiro, A.R., and Brickman, J.M. (2020). From pluripotency to totipotency: an experimentalist’s guide to cellular potency. Development 147. 10.1242/dev.189845.

19. Rossant, J. (2008). Stem Cells and Early Lineage Development. Cell 132, 527–531. 10.1016/j.cell.2008.01.039

20. Macfarlan, T.S., Gifford, W.D., Driscoll, S., Lettieri, K., Rowe, H.M., Bonanomi, D., Firth, A., Singer, O., Trono, D., and Pfaff, S.L. (2012). Embryonic stem cell potency fluctuates with endogenous retrovirus activity. Nature 487, 57–63. 10.1038/nature11244.

21. De Iaco, A., Planet, E., Coluccio, A., Verp, S., Duc, J., and Trono, D. (2017). DUX-family transcription factors regulate zygotic genome activation in placental mammals. Nature Genetics 49, 941–945. 10.1038/ng.3858.

22. Fu, X., Djekidel, M.N., and Zhang, Y. (2020). A transcriptional roadmap for 2C-like–to–pluripotent state transition. Science Advances 6. 10.1126/sciadv.aay5181.

23. Shen, H., Yang, M., Li, S., Zhang, J., Peng, B., Wang, C., Chang, Z., Ong, J., and Du, P. (2021). Mouse totipotent stem cells captured and maintained through spliceosomal repression. Cell 184, 2843–2859.e2820. 10.1016/j.cell.2021.04.020.

24. Yang, M., Yu, H., Yu, X., Liang, S., Hu, Y., Luo, Y., Izsvák, Z., Sun, C., and Wang, J. (2022). Chemical-induced chromatin remodeling reprograms mouse ESCs to totipotent-like stem cells. Cell Stem Cell 29, 400–418.e413. 10.1016/j.stem.2022.01.010.

25. Hu, Y., Yang, Y., Tan, P., Zhang, Y., Han, M., Yu, J., Zhang, X., Jia, Z., Wang, D., Yao, K., et al. (2022). Induction of mouse totipotent stem cells by a defined chemical cocktail. Nature 617, 792–797 10.1038/s41586-022-04967-9.

26. Huang, B., Peng, X., Zhai, X., Hu, J., Chen, J., Yang, S., Huang, Q., Deng, E., Li, H., Barakat, T.S., et al. (2024). Inhibition of HDAC activity directly reprograms murine embryonic stem cells to trophoblast stem cells. Developmental Cell 59, 2101–2117.e2108. 10.1016/j.devcel.2024.05.009.

27. Zhang, P., Zhai, X., Huang, B., Sun, S., Wang, W., and Zhang, M. (2023). Highly efficient generation of blastocyst-like structures from spliceosomes-repressed mouse totipotent blastomere-like cells. Science China Life Sciences 66, 423–435 10.1007/s11427-022-2209-3.

28. Li, R., Zhong, C., Yu, Y., Liu, H., Sakurai, M., Yu, L., Min, Z., Shi, L., Wei, Y., Takahashi, Y., et al. (2019). Generation of Blastocyst-like Structures from Mouse Embryonic and Adult Cell Cultures. Cell 179, 687–702 e618. 10.1016/j.cell.2019.09.029.

29. Zhang, P., Zhai, X., Huang, B., Sun, S., Wang, W., and Zhang, M. (2023). Highly efficient generation of blastocyst-like structures from spliceosomes-repressed mouse totipotent blastomere-like cells. Sci China Life Sci. 10.1007/s11427-022-2209-3.

30. Yamanaka, Y., Lanner, F., and Rossant, J. (2010). FGF signal-dependent segregation of primitive endoderm and epiblast in the mouse blastocyst. Development 137, 715–724. 10.1242/dev.043471.

31. Shen, H., Yang, M., Li, S., Zhang, J., Peng, B., Wang, C., Chang, Z., Ong, J., and Du, P. (2021). Mouse totipotent stem cells captured and maintained through spliceosomal repression. Cell 184, 2843–2859 e2820. 10.1016/j.cell.2021.04.020.

32. Ring, D.B., Johnson, K.W., Henriksen, E.J., Nuss, J.M., Goff, D., Kinnick, T.R., Ma, S.T., Reeder, J.W., Samuels, I., Slabiak, T., et al. (2003). Selective Glycogen Synthase Kinase 3 Inhibitors Potentiate Insulin Activation of Glucose Transport and Utilization In Vitro and In Vivo. Diabetes 52, 588–595. 10.2337/diabetes.52.3.588.

33. Aulicino, F., Pedone, E., Sottile, F., Lluis, F., Marucci, L., and Cosma, M.P. (2020). Canonical Wnt Pathway Controls mESC Self-Renewal Through Inhibition of Spontaneous Differentiation via β-Catenin/TCF/LEF Functions. Stem Cell Reports 15, 646–661. 10.1016/j.stemcr.2020.07.019.

34. Xie, H., Tranguch, S., Jia, X., Zhang, H., Das, S.K., Dey, S.K., Kuo, C.J., and Wang, H. (2008). Inactivation of nuclear Wnt-â-catenin signaling limits blastocyst competency for implantation. Development 135, 717–727. 10.1242/dev.015339.

35. de Vries, W.N., Evsikov, A.V., Haac, B.E., Fancher, K.S., Holbrook, A.E., Kemler, R., Solter, D., and Knowles, B.B. (2004). Maternal β-catenin and E-cadherin in mouse development. Development 131, 4435–4445. 10.1242/dev.01316.

36. Messerschmidt, D., de Vries, W.N., Lorthongpanich, C., Balu, S., Solter, D., and Knowles, B.B. (2016). β-catenin-mediated adhesion is required for successful preimplantation mouse embryo development. Development 143, 1993–1999. 10.1242/dev.133439.

37. Clift, D., So, C., McEwan, W.A., James, L.C., and Schuh, M. (2018). Acute and rapid degradation of endogenous proteins by Trim-Away. Nature Protocols 13, 2149–2175. 10.1038/s41596-018-0028-3.

38. Stepniak, E., Radice, G.L., and Vasioukhin, V. (2009). Adhesive and Signaling Functions of Cadherins and Catenins in Vertebrate Development. Cold Spring Harbor Perspectives in Biology 1, a002949–a002949. 10.1101/cshperspect.a002949.

39. Kobielak, A., and Fuchs, E. (2004). α-catenin: at the junction of intercellular adhesion and actin dynamics. Nature Reviews Molecular Cell Biology 5, 614–625. 10.1038/nrm1433.

40. Harris, T.J.C., and Tepass, U. (2010). Adherens junctions: from molecules to morphogenesis. Nature Reviews Molecular Cell Biology 11, 502–514. 10.1038/nrm2927.

41. Stephenson, R.O., Yamanaka, Y., and Rossant, J. (2010). Disorganized epithelial polarity and excess trophectoderm cell fate in preimplantation embryos lacking E-cadherin. Development 137, 3383–3391. 10.1242/dev.050195.

42. Rivron, N.C., Frias-Aldeguer, J., Vrij, E.J., Boisset, J.C., Korving, J., Vivie, J., Truckenmuller, R.K., van Oudenaarden, A., van Blitterswijk, C.A., and Geijsen, N. (2018). Blastocyst-like structures generated solely from stem cells. Nature 557, 106–111. 10.1038/s41586-018-0051-0.

43. Li, H., Guan, W., Huang, J., Shen, P., Wu, J., Luo, H., Yang, Y., Ning, S., Chang, L., Zhao, H., et al. (2025). A complete model of mouse embryogenesis through organogenesis enabled by chemically induced embryo founder cells. Cell. 10.1016/j.cell.2025.07.018.

44. Liu, K., Yan, Z., Bai, D., Jiang, R., Bi, Y., Ma, X., Xiang, J., Sheng, Y., Dong, B., Ning, Z., et al. (2025). Modeling post-gastrula development via bidirectional pluripotent stem cells. Cell Research 35, 954–969. 10.1038/s41422-025-01172-x.

